# Molecular Dynamics Simulation as a Promising Approach for Computational Study of Liquid Crystal-based Aptasensors

**DOI:** 10.1101/2022.11.19.517204

**Authors:** Hamed Zahraee, Seyed Shahriar Arab, Zahra Khoshbin, Seyed Mohammad Taghdisi, Khalil Abnous

**Author notes:** Corresponding Authors: **Prof. Khalil Abnous** Professor of Department of Medicinal Chemistry Pharmaceutical Research Center, School of Pharmacy Mashhad University of Medical Sciences Mashhad, Iran.: **Email:**, **Dr. Zahra Khoshbin** Pharmaceutical Research Center, Pharmaceutical Technology Institute Mashhad University of Medical Sciences Mashhad, Iran.: **Email:**.

## Abstract

As a potent computational methodology, molecular dynamics (MD) simulation provides advantageous knowledge about biological compounds from the molecular viewpoint. In particular, MD simulation gives exact information about aptamer strands, such as the short synthetic oligomers, their orientation, binding sites, folding-unfolding state, and conformational re-arrangement. Also, the effect of the different chemicals and biochemicals as the components of aptamer-based sensors (aptasensors) on the aptamer-target interaction can be investigated by MD simulation. Liquid crystals (LCs) as soft substances with characteristics of both solid anisotropy and liquid fluidity are new candidates for designing label-free aptasensors. To now, diverse aptasensors have been developed experimentally based on the optical anisotropy, fluidity, and long-range orientational order of LCs. Here, we represent a computational model of an LC-based aptasensor through a detailed MD simulation study. The different parameters are defined and studied to achieve a comprehensive understanding of the computational design of the LC-based aptasensor, including the density of LCs, their orientation angle, and lognormal distribution in the absence and presence of aptamer strands, both aptamer and target molecules with various concentrations, and interfering substance. As a case study, the tobramycin antibiotic is considered the target molecule for the computational model of the LC-based aptasensor.

## 1. Introduction

Biosensors have gained significant attention as highly efficient analytical devices in various fields of science, such as healthcare, pharmaceuticals, clinical diagnostics, environmental monitoring, and food quality control (Azzouz et al., 2022; Khoshbin et al., 2023; Zavvar et al., 2022). Generally, biosensors can include diverse types of bioreceptor segments for the specific targeting of molecules, such as antibodies, peptides, nucleic acids, cells, enzymes, aptamers, and more. Among these, biosensors based on aptamers, known as aptasensors, are some of the most effective and well-organized biosensing tools available (An et al., 2022; Du & Dong, 2017; Hamed Zahraee, Atiyeh Mehrzad, et al., 2023). Aptamers are synthetic single-stranded oligonucleotides that exhibit high binding affinity to a wide range of targets and are obtained through the systematic evolution of ligands by the exponential enrichment (SELEX) selection method (Darmostuk, Rimpelova, Gbelcova, & Ruml, 2015; Ellington & Szostak, 1990). This in-vitro production method eliminates the need for laboratory animals to synthesize aptamers. Moreover, the small size and low weight of aptamers makes their generation and modification facile and cost-effective (Hamed Zahraee, Zahra Khoshbin, et al., 2023). Aptamers possess long shelf life, less immunogenicity and toxicity, great stability, biocompatibility, and reversible thermal denaturation (Keefe, Pai, & Ellington, 2010). These outstanding advantages make aptamers efficacious receptors for diverse biological sensing approaches. Additionally, aptasensors have a fast response time, low cost, and great target detection limit, making them user-friendly analytical tools (Khoshbin, Verdian, Housaindokht, Izadyar, & Rouhbakhsh, 2018; Zhu et al., 2022).

Liquid crystals (LCs) are unique materials composed of aromatic molecules with side chains that exhibit a state between isotropic liquid and crystalline solid (Pilz da Cunha, Debije, & Schenning, 2020). Due to their long-range orientational order, responsiveness to external stimuli, elastic strain, optical anisotropy, and susceptibility to orientation, LCs have become an attractive choice for biosensor design (Devi, Pani, & Pal, 2022; Zahra Khoshbin, Khalil Abnous, Seyed Mohammad Taghdisi, & Asma Verdian, 2021; Pani, Sil, & Pal, 2023). LC-based biosensors are a promising candidate for label-free and portable target monitoring assays, as they enable the detection of targets through visualizing the orientation of the LCs using a polarized optical microscope, without the need for labeled probes that can be costly and time-consuming (Hussain, Pina, & Roque, 2009; Jang et al., 2005; Verdian, Rouhbakhsh, & Fooladi, 2021; Y. Wang, Hu, Tian, Gao, & Yu, 2016).

Molecular dynamics (MD) simulation is a potent computational methodology that provides valuable knowledge of biological components (M.-R. Li, Zhang, & Zhang, 2018; Monti, Cappelli, Bronco, Giusti, & Ciardelli, 2006; Nicholls et al., 2009; Plattner, Doerr, De Fabritiis, & Noe, 2017; Zahraee, Arab, Khoshbin, & Bozorgmehr, 2023). By simulating the molecular interactions in a biological system, MD simulation can provide accurate information about binding sites, structural rearrangements, and the folding-unfolding status of aptamers (Bini, Mascini, Mascini, & Turner, 2011; Gao et al., 2016; Kim, Rhee, & Shin, 2018; Luo & Mu, 2016; Mary, Haris, Varghese, Aparna, & Sudarsanakumar, 2017; Phanchai, Srikulwong, Chompoosor, Sakonsinsiri, & Puangmali, 2018). Additionally, MD simulation can be used to study the dynamics, structure, and energetics of biological systems under a range of experimental conditions, including pressure, pH, and temperature (Aho et al., 2022; Johnson, Kohlmeyer, Johnson, & Klein, 2009; Moradi, Shareghi, Saboury, & Farhadian, 2020; Pal, Aich, Chakraborty, & Jana, 2022). MD simulation can complement experimental methods by preventing incorrect interpretations of experimental results (Cao et al., 2017; Khanniche et al., 2017; Kutnowski et al., 2018; Zahraee, Mohammadi, Parvaee, Khoshbin, & Arab, 2023; Zeng et al., 2016). Moreover, it can provide basic information at the atomistic level when experimental assays have limitations or are incapable (Nada & Furukawa, 2012; Rigoldi et al., 2016; Shao et al., 2021; J. Wang, Qian, Li, & Qiu, 2020).

In this study, we emphasize the potency of MD simulation as a complement to experimental assays in designing an LC-based aptasensor. Specifically, we present a computational model of an LC-based aptasensor through a detailed MD simulation study. As a case study, we selected the tobramycin antibiotic as the target molecule. Tobramycin as an anti-infective antibiotic drug commonly utilizes for therapeutic and prophylactic purposes, because of its good water-solubility, low cost, and antimicrobial effects (DiCicco, Duong, Chu, & Jansen, 2003; Rosalia et al., 2022). However, its abuse can cause adverse effects in human bodies, such as neuromuscular blocking, nephrotoxicity, and hypersensitivity (Ma, Wang, Jia, & Xiang, 2018; Shang et al., 2019; S. Wang et al., 2019). To date, several strategies have been applied to detect tobramycin, including high-performance liquid chromatography, UV-Visible absorption spectroscopy, liquid chromatography-mass spectrometry, and capillary electrophoresis (Blanchaert, Huang, Wach, Adams, & Van Schepdael, 2017; Law et al., 2006; Meng et al., 2021; Mukhtar, Mamat, & See, 2018). Since the methods are time-consuming, high-cost, and operator-dependent, the LC-based aptasensors have been developed for target detection (D. Li et al., 2022; Zhang, Li, Han, Yuan, & He, 2022). However, LC-based aptasensors are impaired by high temperatures and light, leading to a loss of the LCs’ contrast (O’Neill & Kelly, 2003; Pagano-Stauffer, Johnson, Clark, & Handschy, 1986). Hence, we emphasize the potency of MD simulation as a complement to experimental assays in studying an LC-based aptasensor. Indeed, a MD simulation study of LC-based aptasensors can reduce the cost and time of experimental research.

## 2. Computational Details

The crystal structure of tobramycin and specific aptamer was taken from the protein data bank (PDB ID: 1LC4) to construct the MD input files. The sequence of 5’-ACT TGG TTT AGG TAA TGA GTA-3’ was obtained through nucleotide exchange by YASARA software (Krieger & Vriend, 2014) and modification of hydrogen deficiency using PyMOL software (The PyMOL Molecular Graphics System, Version 1.7.2.1 Schrödinger, LLC.) (DeLano, 2002). Since the orientation of LCs can be easily disrupted by aptamer sequences with more than 24 base units (Z. Khoshbin, K. Abnous, S. M. Taghdisi, & A. Verdian, 2021), two adenosine (A) nucleotides were added to the 3’ end of the aptamer as the spacer. The MD simulations were carried out by GROMACS 2020.2 (Abraham et al., 2015) with the standard CHARMM27 force field (Brooks et al., 2009; Patel, Mackerell, & Brooks, 2004). The structure of 4-Cyano-4’-pentylbiphenyl (5CB) LC was drawn by ChemDraw software (Z. Li, Wan, Shi, & Ouyang, 2004). Its structural and energetic optimization was done by Chem3D (Z. Li et al., 2004) and Gaussian 09 software (Frisch, 2009) by DFT (B3LYP) method within the 6-31++G(2d,2p) basis set. The topology parameters for tobramycin and LC were obtained by using SwissParam server (Zoete, Cuendet, Grosdidier, & Michielin, 2011). Six rectangular boxes were designed in which the aptamers were fixed on their bottom. One system was considered as the control with no tobramycin molecule. Another box was considered for the interfering system containing 5 gentamycin molecules as the interfering compound. Four others included 1, 5, 20, and 40 tobramycin molecules. The boxes were filled with 450 LC molecules. Charge neutrality was done by adding an adequate number of Na^+^ and Cl^-^ ions to the boxes. The details of the simulated systems is given in Table 1. Simulations were carried out using a cut-off distance of 1.4 nm with a time step of 1 fs. The steepest-descent algorithm (Fletcher & Powell, 1963) was applied to minimize the energy of the system. The velocity rescaling thermostat and Parrinello-Rahman barostat were applied for the temperature and pressure coupling, respectively (Bussi, Donadio, & Parrinello, 2007; Parrinello & Rahman, 1981). Periodic boundary conditions (PBC) were introduced in all directions to implement the MD simulations. Finally, 180 ns production MD simulation of each system was performed with a time step of 1 fs at 300 K. The Particle Mesh Ewald (PME) method was considered to obtain the long-range electrostatic interactions (Darden, York, & Pedersen, 1993).

**Table 1.**
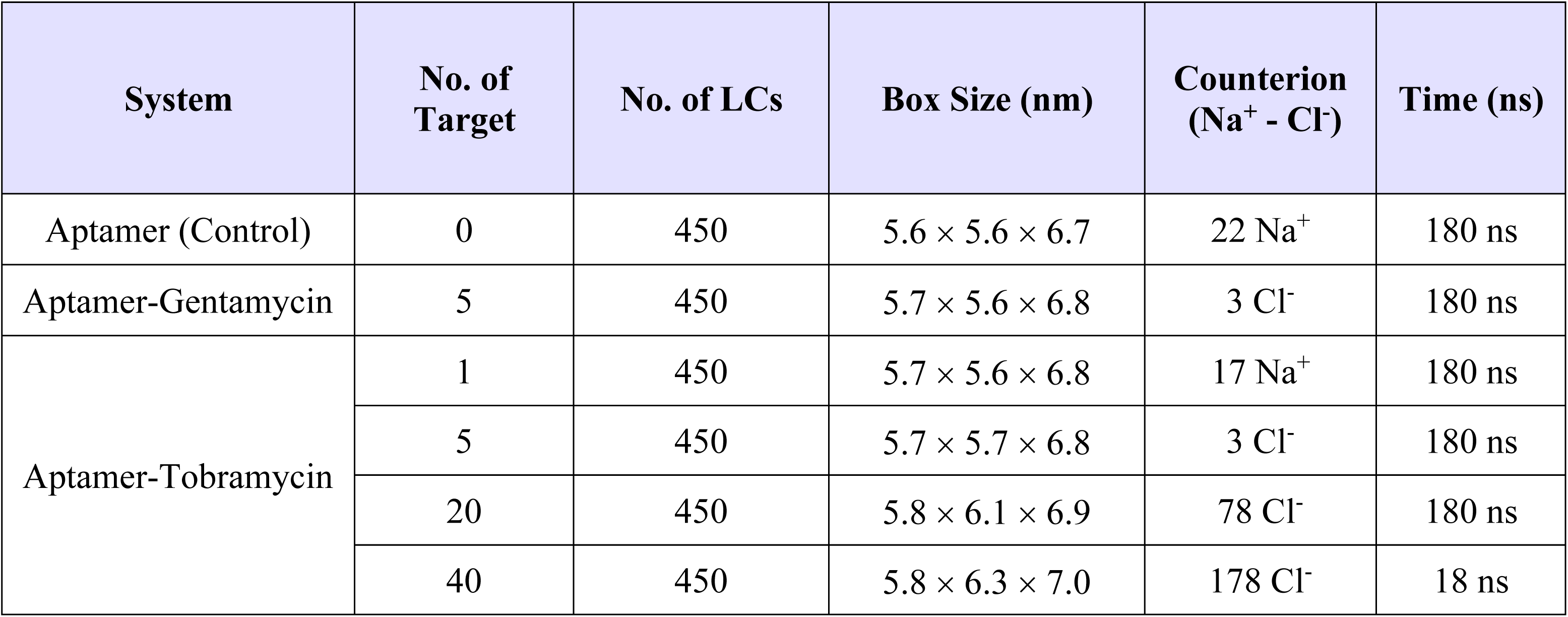
Details of the simulated systems.

| System | No. of Target | No. of LCs | Box Size (nm) | Counterion (Na<sup>+</sup> - Cl<sup>-</sup>) | Time (ns) |
| --- | --- | --- | --- | --- | --- |
| Aptamer (Control) | 0 | 450 | 5.6 × 5.6 × 6.7 | 22 Na<sup>+</sup> | 180 ns |
| Aptamer-Gentamycin | 5 | 450 | 5.7 × 5.6 × 6.8 | 3 Cl<sup>-</sup> | 180 ns |
| Aptamer-Tobramycin | 1 | 450 | 5.7 × 5.6 × 6.8 | 17 Na<sup>+</sup> | 180 ns |
|  | 5 | 450 | 5.7 × 5.7 × 6.8 | 3 Cl<sup>-</sup> | 180 ns |
|  | 20 | 450 | 5.8 × 6.1 × 6.9 | 78 Cl<sup>-</sup> | 180 ns |
|  | 40 | 450 | 5.8 × 6.3 × 7.0 | 178 Cl<sup>-</sup> | 18 ns |

## 3. Results and Discussions

To introduce MD simulation as a potent approach for computational model of LC-based aptasensor, six different systems were designed and studied by MD simulation, including 1) control system containing fixed tobramycin aptamer at the bottom of simulation box and LCs, 2) interfering system containing fixed tobramycin aptamer, LCs, and 5 gentamycin molecules, 3) target systems containing fixed aptamer, LCs, and tobramycin molecules with the number of 1, 5, 20, and 40 (named A1T, A5T, A20T, and A40T, respectively).

The structural stability of the studied systems was investigated by analyzing the root mean square deviation (RMSD). Figure 1A indicates the RMSD plots of the simulated systems, proving that they were in the equilibration state after the MD simulation. Moreover, 180 ns was considered as the proper time for having precise results from further analyses. The results clarify the greater stability of all the target systems in comparison with both the control and interfering systems.

**Figure 1.**
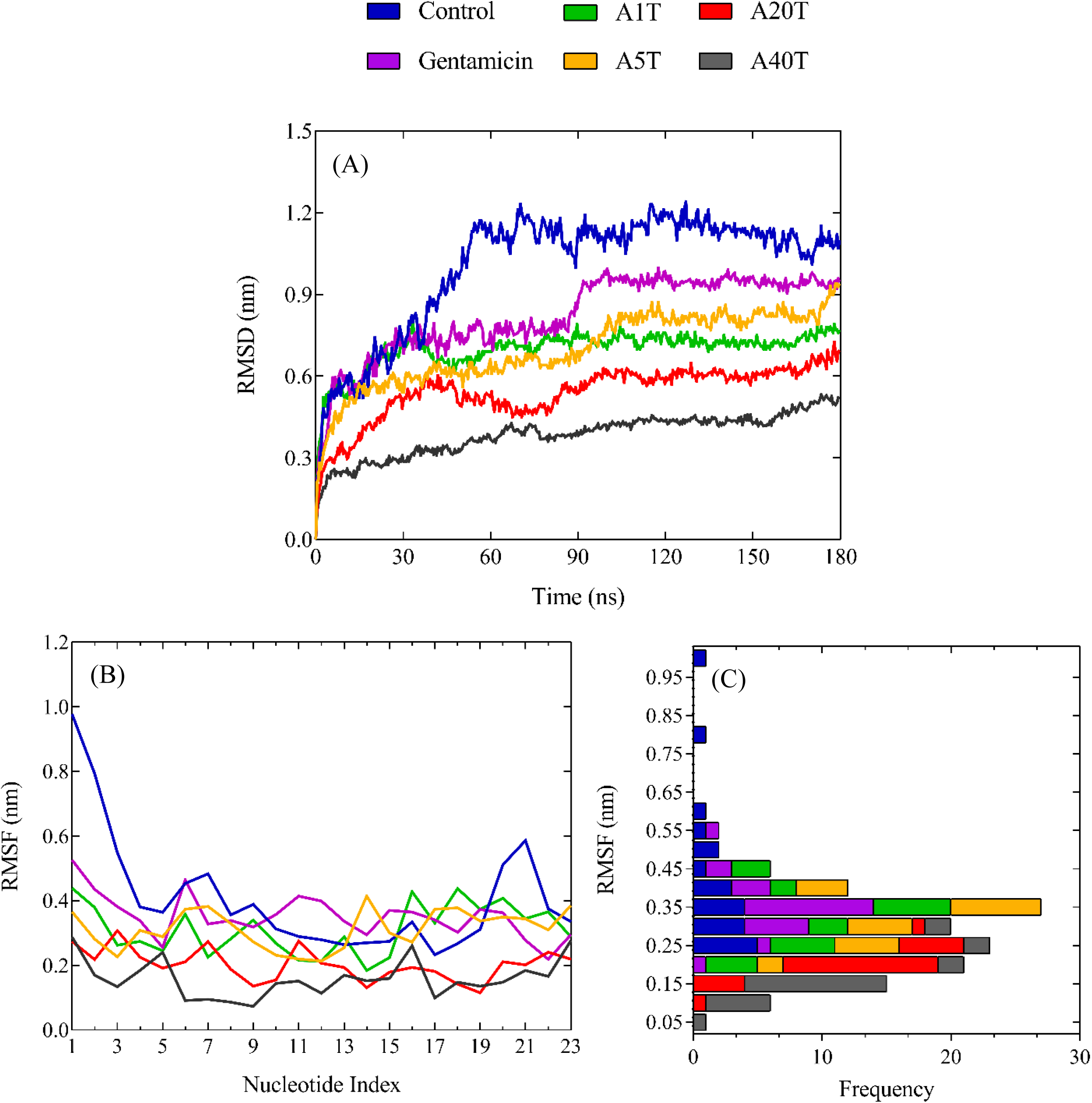
RMSD, RMSF, and RMSF frequency analyses (A-C) for the simulated systems during 180 ns MD simulation.

Root mean square fluctuation (RMSF) is a criterion for conformational flexibility of DNA strands over the simulation time (Chakraborty, Khatua, & Bandyopadhyay, 2016; M. Li et al., 2013). The RMSF plot for the aptamer in the absence and presence of the different numbers of tobramycin, and also, in the presence of interfering compound was depicted against the residue numbers during the MD simulation time in Figure 1B. As it is obvious, the aptamer flexibility decreased in some residues with increasing the number of tobramycin molecules in the systems, which may be due to more interaction of the aptamer with the target molecules. Generally, the middle nucleotides of the aptamer possessed less flexibility than the terminal ones in the systems including targets. Figure 1C indicates the frequency of RMSF for the simulated systems in the range of 0.15-0.45 nm.

Figure 2 indicates the situation of the LC molecules in the different simulated systems in the last frame of the MD simulation. As shown in Figure 2, LCs were in the same direction with the vertical alignment in the control system free of tobramycin, proving the capability of the aptamer strand with nucleotides lower than 24 to regularly align LCs.

**Figure 2.**
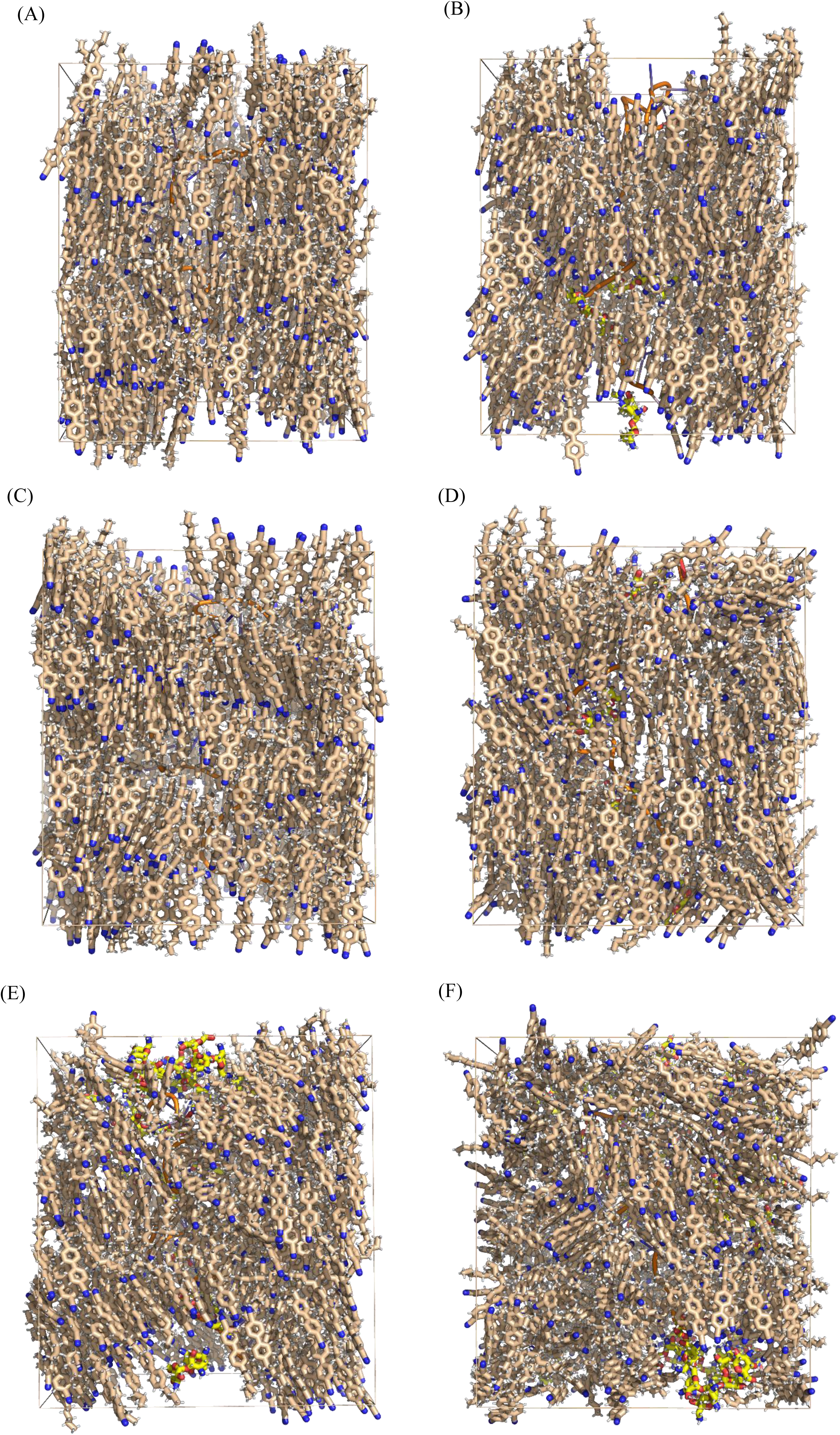
The situation of the LC components of the simulated systems in the last frame of the MD simulation. (A) control system free of tobramycin, (B) system containing gentamycin as the interfering compound, (C-F) systems with 1, 5, 20, and 40 tobramycin molecules, respectively (the phosphate chain of the aptamer is shown to be orange, and the molecule of tobramycin is indicated to be yellow in color).

The diffusion coefficient (D) parameter can reflect the flexibility of the aptamer that affects the regularity of LCs during the MD simulation. Figure S1 illustrates the remarkable difference in the D parameter of the systems in the absence and presence of tobramycin. According to Figure S1, the D parameter in the control and interfering systems was less than the others, highlighting the less flexibility of the aptamer, and therefore, vertical alignment of LCs. Proximity of the D parameter of interfering system to the control system clarifies a negligible flexibility of the aptamer through no interaction with gentamycin molecules, and therefore, vertical alignment of LCs in this system. The results indicate that D parameter for the aptamer enhanced with increasing the number of tobramycin molecules in the studied systems. It proves an increase in the aptamer flexibility with increasing the number of tobramycin molecules. Thus, an increased value of the D index proved the greater irregularity in the vertical orientation of LCs.

To have a detailed insight into the state of the LC molecules for each studied system, their angle changes with respect to the Z axis of the simulation box were investigated. For this purpose, a vertical vector was defined as a basis for the initial state of the LC molecules by considering their C12 and N atoms of the nitrile group (Figure 3A); and its angle with the Z axis of the simulation box. Figure 3B indicates the average of the angle of LCs during 180 ns. It is obvious that the changes in the LCs’ angle were the least for the control system and system containing gentamycin. Besides, the angle changes for these two systems were the same at the last times of the simulation, proving the negligible disruption of the LCs’ alignment in the presence of gentamycin, and also, insignificant interaction of gentamycin with the aptamer. A comparison between the control system and A1T proved that the aptamer-target interaction disturbed the orientation of LC molecules. However, increasing the number of tobramycin molecules to 5 (A5T system) had no considerable effect on the orientation of LCs compared to the A1T system. This confirms that the aptamer binds to tobramycin in a 1:1 stoichiometry, as previously determined in other studies (Vicens & Westhof, 2002; Y. Wang, Killian, Hamasaki, & Rando, 1996). Moreover, an increase in the number of tobramycin molecules to 20 and 40 changed the orientation of LCs. Since the binding stoichiometry of the aptamer to tobramycin was 1:1, the more disruption in the LCs’ orientation in the A20T and A40 systems in comparison with the A1T system was due to the free tobramycin molecules. Having difference with this study, in the experimentally-constructed LC-based aptasensors, the free target molecules are eliminated from the aptasensing system during the washing process, and after that, LCs are injected onto the aptasensing substrate. Hence, what causes the change in the vertical arrangement of LCs in the experimental LC aptasensors is only the target molecules interacted with the aptamer strands. Our theoretical results highlight the importance of washing process in the experimentally-constructed LC-based aptasensors to eliminate the undesirable effect of free targets on the order of LCs. The result clarifies that MD simulation can be complementary to experimental assays for designing LC-based aptasensors.

**Figure 3.**
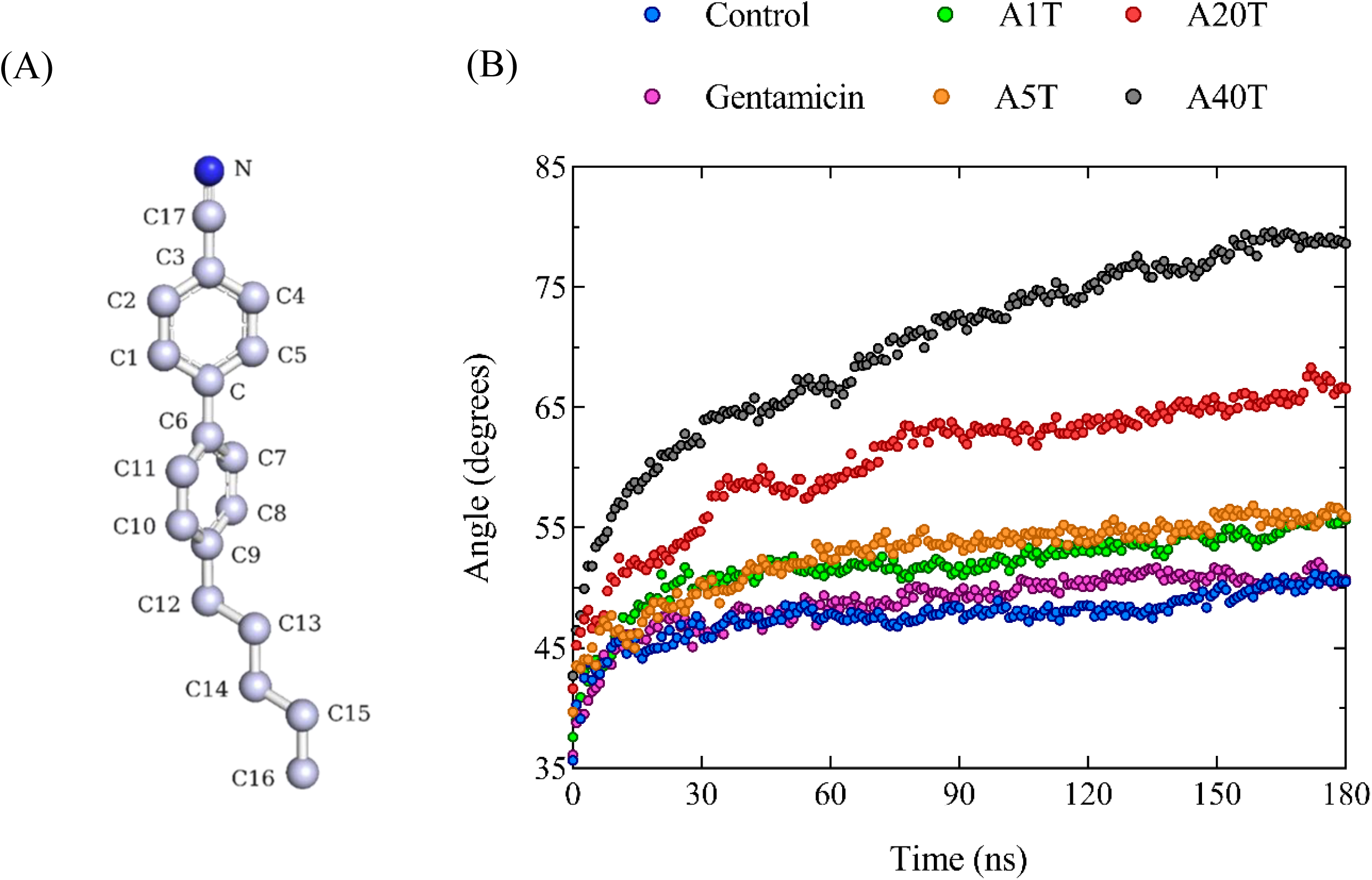
(A) Molecular structure of 4-Cyano-4’-pentylbiphenyl liquid crystal. (B) The average of the angles of LC molecules with respect to the Z axis of the simulation box during 180 ns.

Figure 4 shows the average of the LCs’ angle respect to the Z axis for each LC molecule in the simulated systems during the MD simulation. According to Figure 4, in the control system, most of LC molecules possessed an angle of 30° and some of them had a rotation of almost 180° compared to the Z axis, proving their relatively vertical direction. A similar result was obtained for the interfering system, which confirmed no interaction of gentamycin molecule with the aptamer, and consequently, its high specificity toward tobramycin. In A1T system, the number of LCs with the angle more than 60° increased compared to the control system, indicating the specific interaction of the aptamer with tobramycin molecule. With increasing the number of tobramycin molecules, the angle of LCs enhanced from 30° to the range of 60-120°, which showed the disruption of LCs’ regularity caused by free tobramycin molecules based on 1:1 aptamer-tobramycin binding stoichiometry. For simplification in a display, the angle of LCs at the first, middle, and end of the simulation time was given in Figure S2, which clearly shows the increase in the changes of the LCs’ angle by enhancing the number of tobramycin molecules in the system.

**Figure 4.**
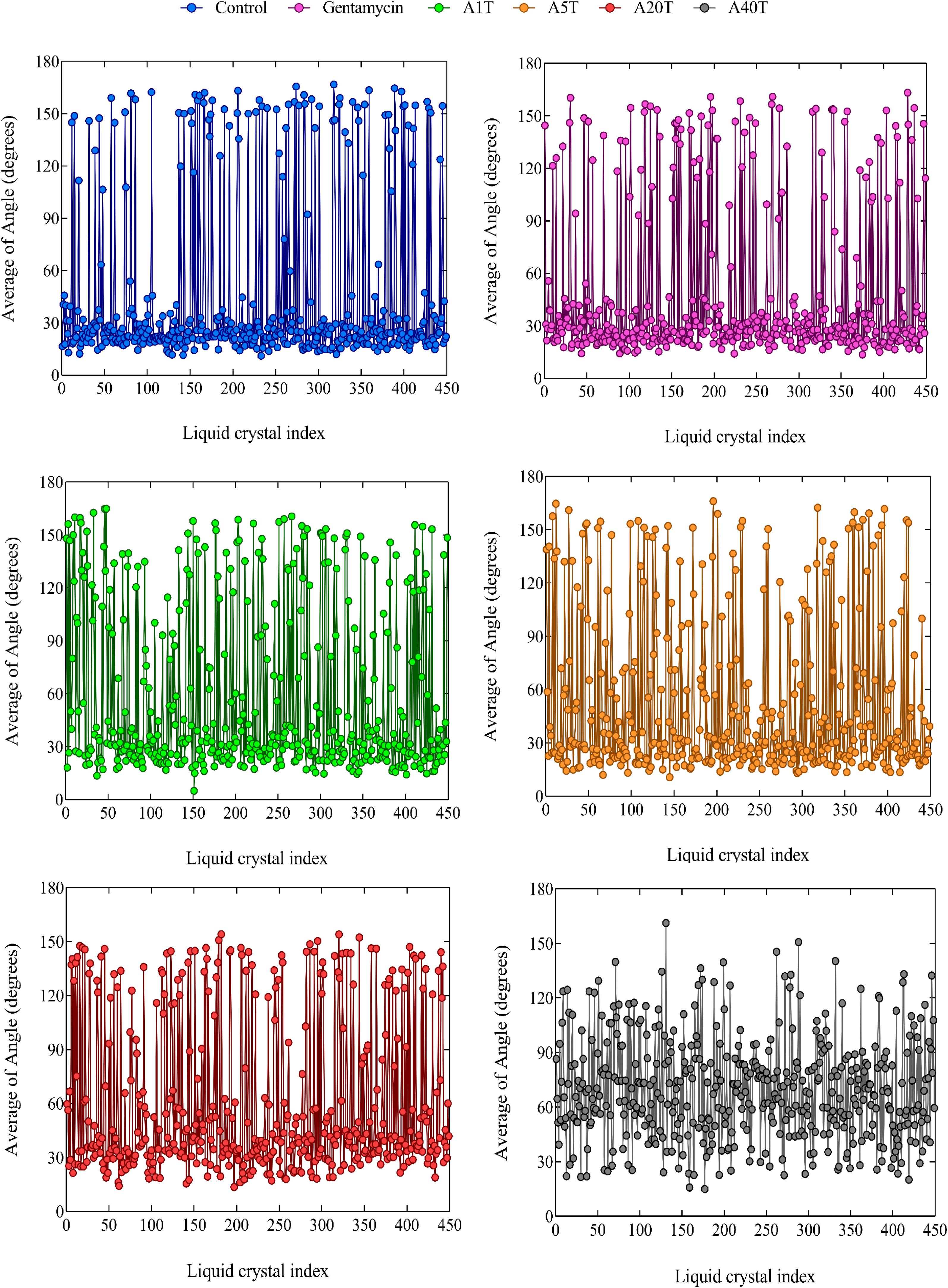
The average of the LCs’ angle with respect to the Z axis of the simulation box for each LC molecule in the simulated systems during the MD simulation.

For further confirmation of the accuracy of the results, the lognormal distribution (Kissell & Poserina, 2017) as a continuous probability distribution of the data was applied based on the following equation:

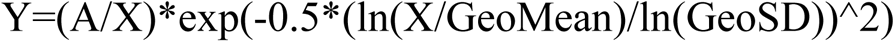

in which Y and X are the probability and angle, respectively. A is related to the amplitude and area of the distribution. GeoMean and GeoSD refer to the geometric mean and geometric standard deviation factor, respectively. Table S1 represents the details and results of the lognormal distribution for the studied systems.

Figure 5 indicates the angle probability of LC molecules along with the lognormal distribution during the MD simulation. The results highlight that the maximum probability was less than 30° in the control system and system containing gentamycin, clarifying the slight deviation of LCs from the vertical direction. With adding tobramycin to the system, the probability partly shifted to 30° due to the aptamer-target interaction. With increasing the number of tobramycin molecules from 1 to 40, a considerable shift in the angle probability to near 90° was achieved, highlighting the change in the direction of LCs from vertical close to horizontal. Also, there was a little probability for the angles in the range of 160°-180°, which was related to the LC molecules with a rotation of almost 180° compared to the Z axis. The results of the lognormal distribution were very close to the probability obtained from the MD output, proving the reliability of the MD results. Moreover, the R-squared values from the lognormal analysis were a confirmation for the accuracy of the MD study.

**Figure 5.**
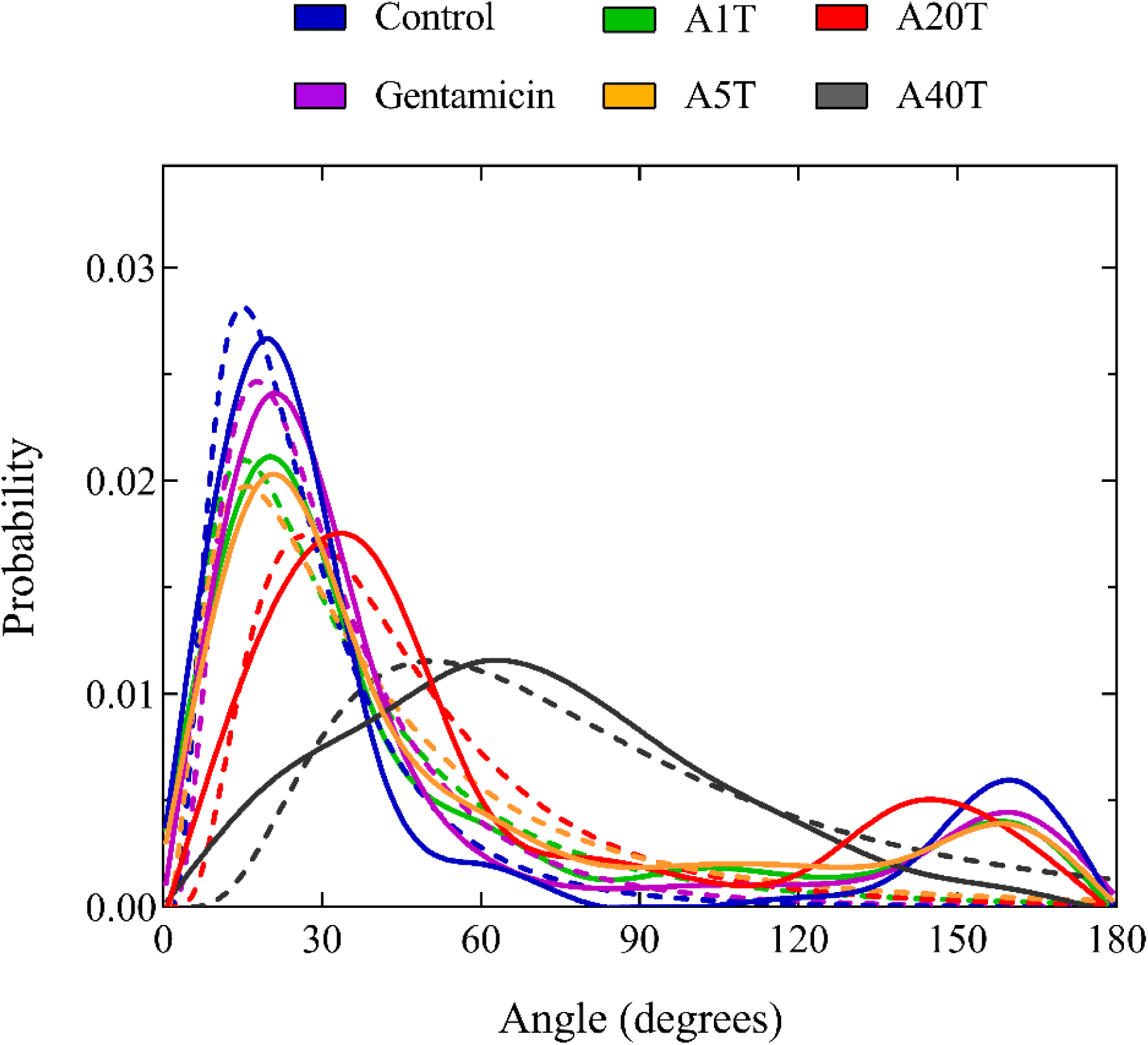
The angle probability of LC molecules was obtained by the lognormal distribution (dashed line) and MD simulation (solid line).

Figure 6 indicates the density of LCs along the X axis in the simulated systems at the initial (0-10 ns), middle (85-95 ns), and end (170-180 ns) of the simulation time. Obviously, the density of LCs changed with passing of the simulation time. For the control and interfering systems, the changes in the density were almost similar; and slightly increased at the end of the MD simulation. In the systems containing tobramycin molecule, the changes in the density of LCs were significant during the MD simulation. An increase in the density of LCs was observed at the end of the MD simulation compared to that for the control system. A comparison between the density of LCs in A1T, A5T, A20T, and A40T systems highlights the significant perturbation in the LCs’ alignment in the presence of more numbers of the target.

**Figure 6.**
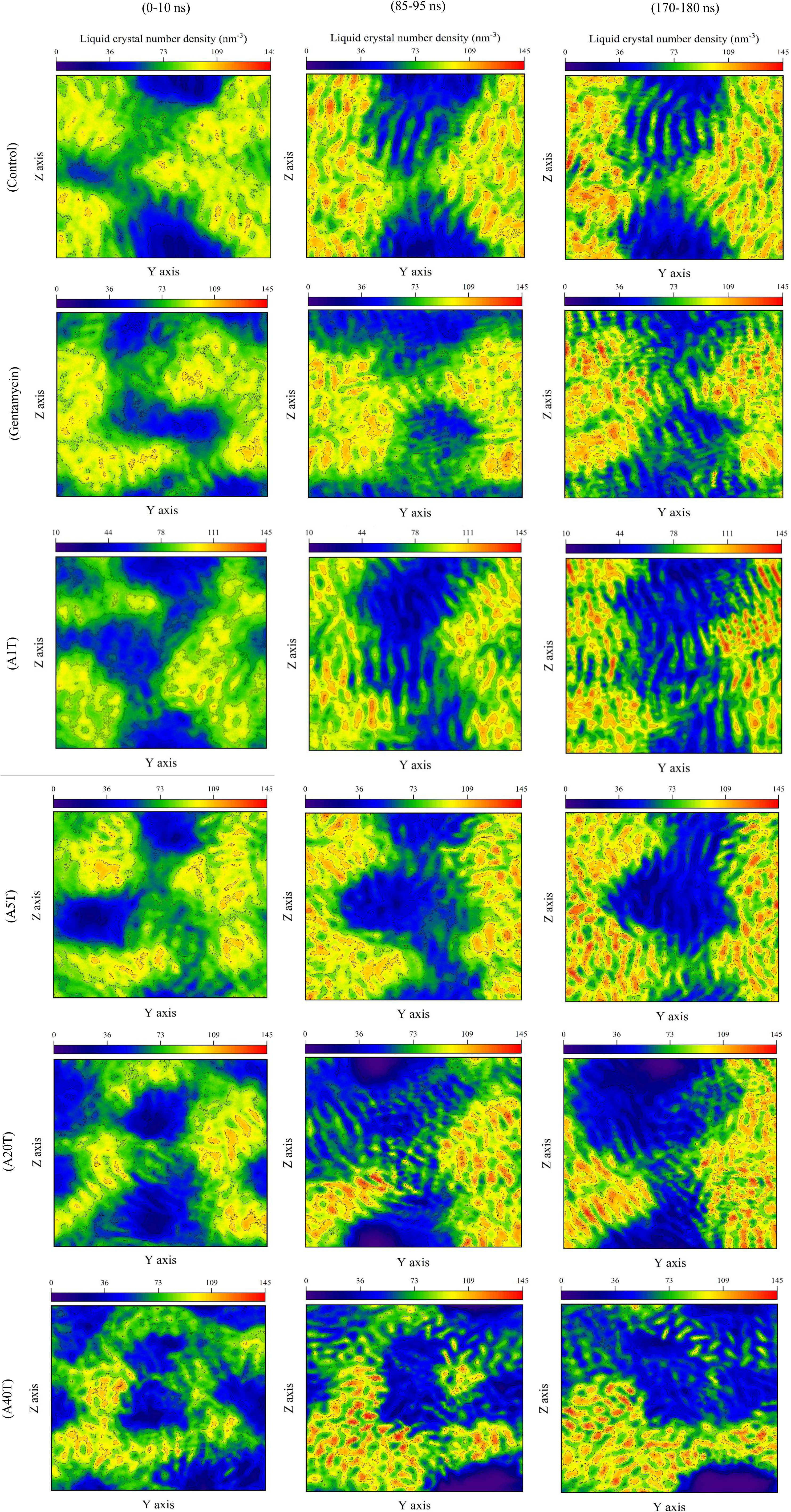
The density (nm^-3^) graph of the LC molecules along the X axis in the different simulated systems at the initial, middle, and end of the simulation time.

The Radial distribution function (RDF) analysis for the studied systems was obtained between the atoms of LCs. Figure S3 indicates that the distance of the C16-C16 atoms of LCs enhanced with increasing the number of the target molecules, due to the more rotation of LCs and greater perturbation of their vertical direction. A comparison with the control system displays that the distance of C16-N atoms of LCs decreased with increasing the number of tobramycin molecules, because of the higher rotation of LC molecules from the vertical state. Finally, the distance between the aptamer nucleotides and tobramycin molecules was calculated for the last 10 ns of the MD simulation (Figure 7). The distance between the target molecule and aptamer nucleotides can be a criterion for the effective interactions between them. In A1T system, the aptamer nucleotides with the index of 7-14 possessed less distance from tobramycin, and therefore, more effective interaction with it. Hence, their distance from the target molecule (1.2 nm and less than that) was chosen as a criterion for specifying the effective binding sites of the aptamer for tobramycin. In A5T system, tobramycin molecules with the index of 1, 4, and 5 had greater interaction with the middle nucleotides of the aptamer, and those with the index of 2 and 3 possessed more effective interactions with the terminal ones. However, only tobramycin with the index of 4 had a distance of 1.2 nm and less from the nucleotides of 7-14 similar to that for A1T system. The result was in accordance with the previous ones corresponding to the 1:1 aptamer-tobramycin binding stoichiometry. For A20T and A40T systems, some tobramycin molecules possessed a distance of 1.2 nm and less from the aptamer nucleotides. However, tobramycin molecules with the index of 1 and 37 in A20T and A40T systems, respectively, occupied the nucleotides with the index of 7-14.

**Figure 7.**
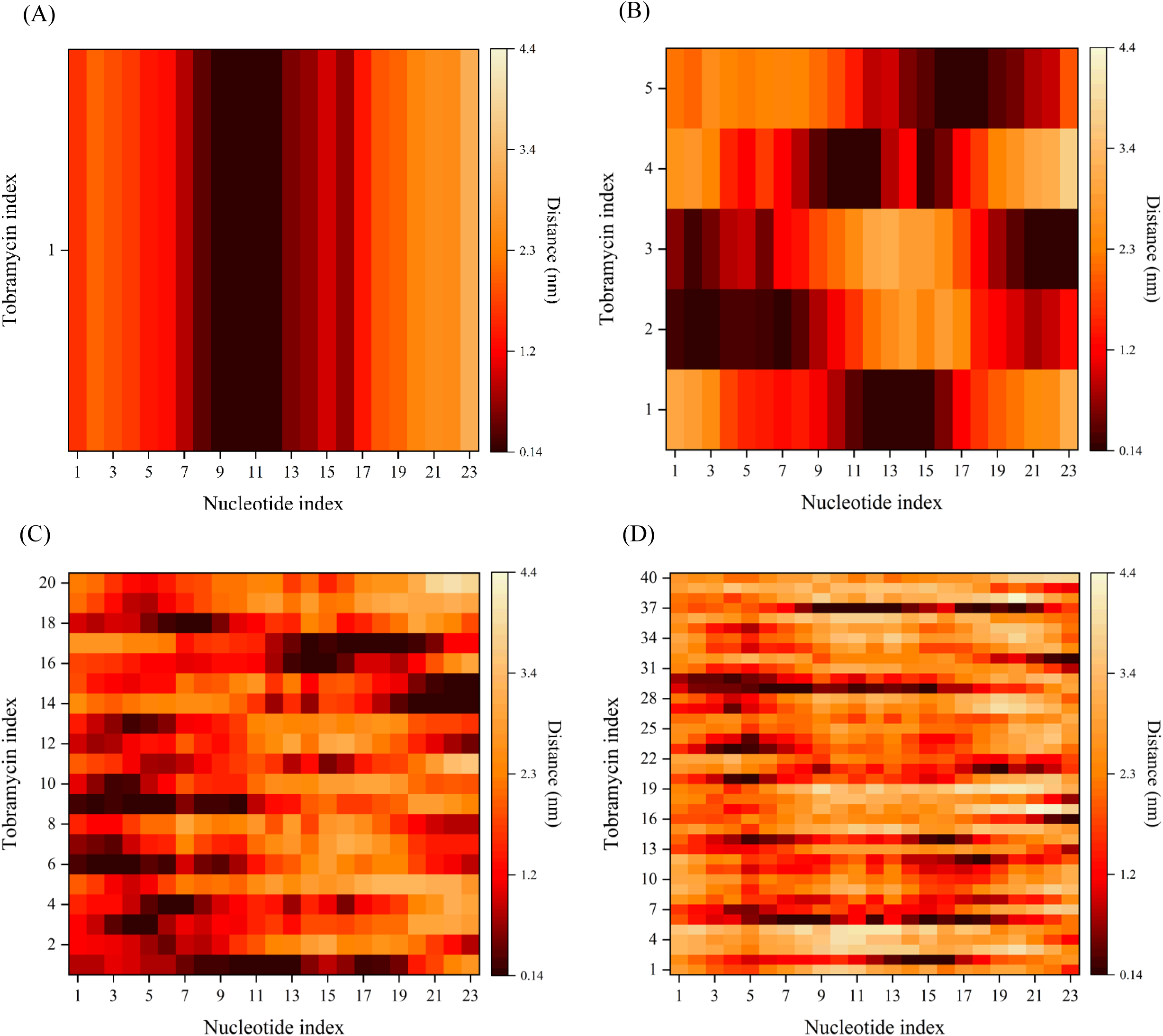
The distance between the tobramycin molecules and the aptamer nucleotides in the A1T, A5T, A20T, and A40T systems (A-D).

Consequently, the aptamer nucleotides with the index of 7-14 could be specified as the binding sites of tobramycin.

## 4. Conclusion

In this research, we studied the potency of MD simulation to computationally model an LC-based aptasensor for the first time. As a case study, the tobramycin antibiotic has been chosen as the target molecule. 180 ns MD simulation was done for the control system, interfering system having gentamycin, and four systems with the different concentrations of tobramycin (A1T, A5T, A20T, and A40T). RMSD results proved the structural stability of the simulated systems after the MD simulation. RMSF analysis indicated that the aptamer flexibility decreased in some residues with increasing the number of tobramycin, which may be due to more interaction of the aptamer with the target molecules. The basis of the LC-aptasensor is change of the vertical alignment of LCs to random in the presence of the target, due to the aptamer-target interaction. Besides, the more aptamer-target complexes, the higher disruption of the vertical arrangement of LCs is. The visualization of the last frame of the simulated systems by PyMOL software indicated that LCs were in the same direction with the vertical alignment in the control system free of tobramycin as well as the interfering system; while the presence of tobramycin molecule significantly disrupted the vertical regularity of LCs. The observations were in accordance with the experimental results (Zahra Khoshbin et al., 2021; Rouhbakhsh, Verdian, & Rajabzadeh, 2020). A detailed insight was achieved by investigating the average of the angle of LCs during 180 ns. The changes in the LCs’ angle were negligible in the control and interfering systems, proving an insignificant interaction of gentamycin with the aptamer. The average of the angle increased in A1T system that confirmed the significant changes in the vertical alignment of LCs due to the great specific interaction of the aptamer with the target. With increasing the number of tobramycin molecules from 1 to 40, a great shift in the angle probability to near 90° was achieved, proving the change in the direction of LCs from vertical close to horizontal. Based on 1:1 aptamer-tobramycin binding stoichiometry, the disruption of LCs’ regularity in A5T, A20T, and A40T systems was induced by free tobramycin molecules. The density of LCs in the systems containing tobramycin remarkably changed during the MD simulation. A comparison between the density of LCs in A1T, A5T, A20T, and A40T systems proved the significant irregularity in the LCs’ alignment in the presence of more numbers of the target. The study confirmed the potency of MD simulation for study of the LC including biosystems. Besides, LC-based aptasensors can be computationally designed by MD simulation and establishing a relationship between the MD-output parameters, such as the density of LCs and their angle change, with the concentration of the desired target.

## Acknowledgment

Financial support for this study was provided by Mashhad University of Medical Sciences. We also thank Mr. Ali Akbar Bromideh (Faculty of Mathematical Sciences, Shahid Beheshti University, Tehran, Iran) for providing statistical support.

---

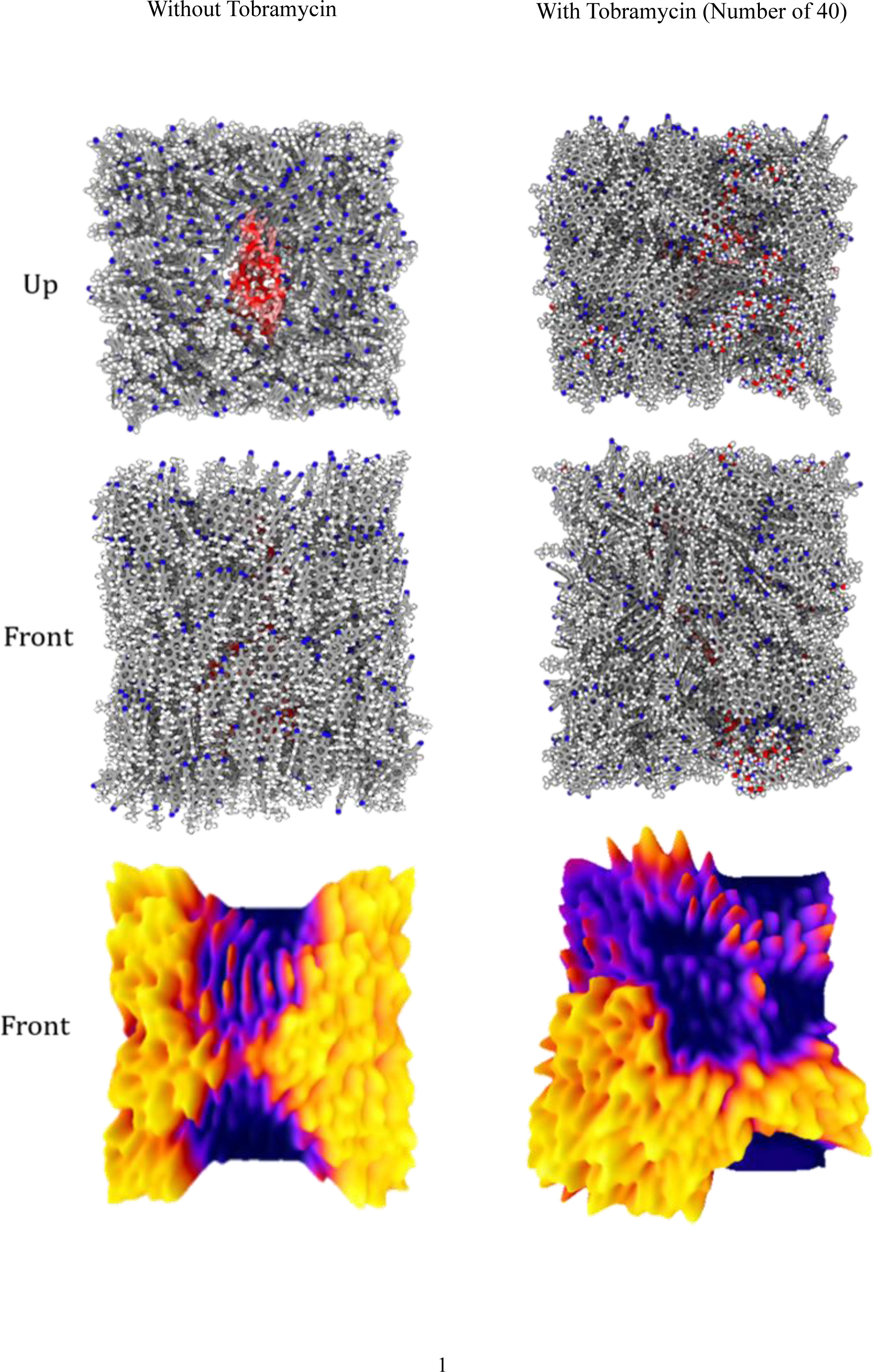

