## Supplementary Material for "Molecular Dynamics Simulation as a Promising Approach for Computational Study of Liquid Crystal-based Aptasensors"

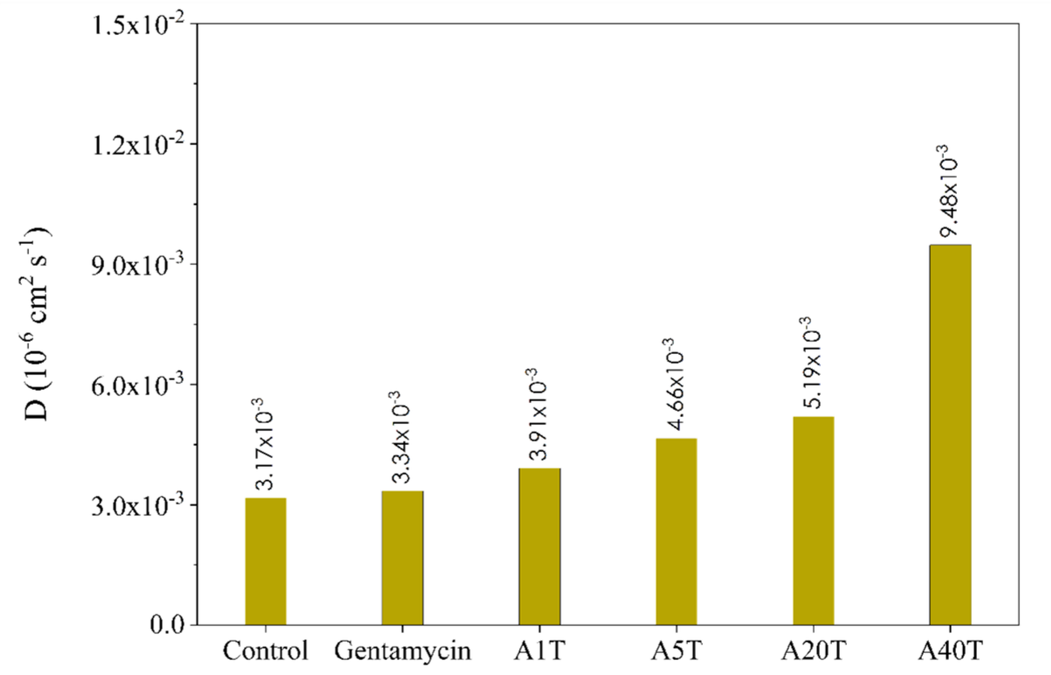


**Figure S1.** The D parameter plot for the aptamer molecule in different simulated systems for the last 20 ns of MD simulation.

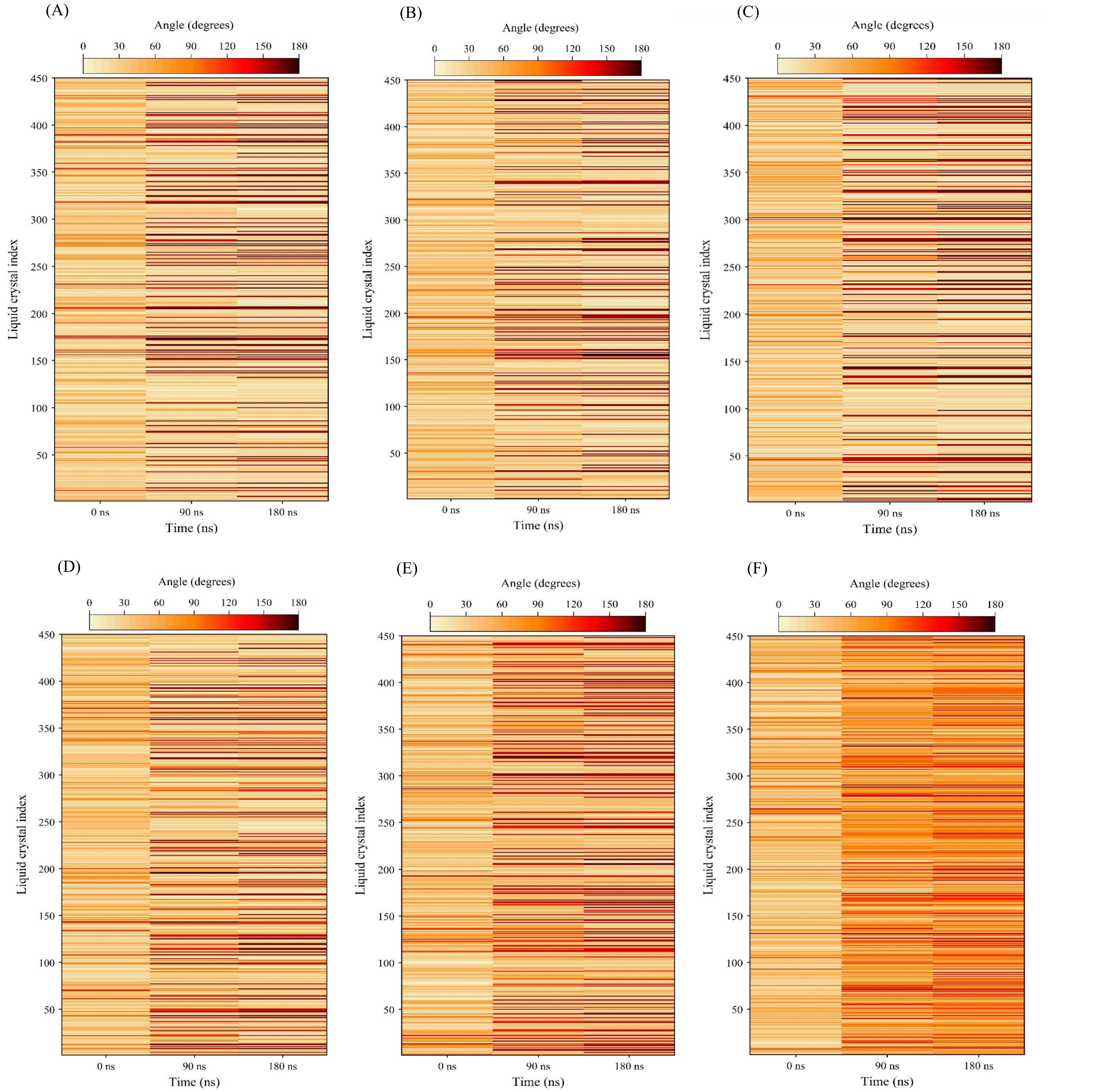


**Figure S2.** The angle of LCs included in the systems at the first, middle, and end of the simulation time; (A) control system, (B) interfering system, (C) A1T, (D) A5T, (E) A20T, and (F) A40T.

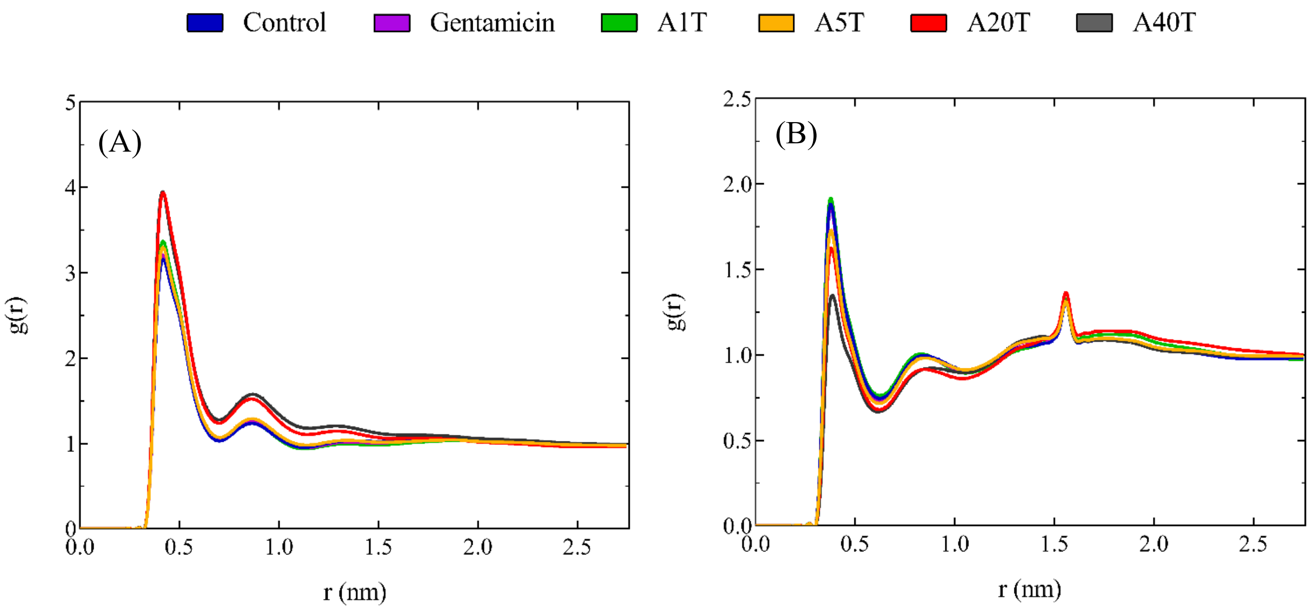


**Figure S3.** The RDF graphs between the C16-C16 of LC molecule (A), and C16-N atoms of LC (B).

**Table S1.** The details of lognormal distribution and the obtained results for the simulated systems.

| Systems | Number of points (X and Y values)* | GeoMean | GeoSD | LnGeoMean | LnGeoSD | Degrees of Freedom | R squared | Sum of Squares |
|---|---|---|---|---|---|---|---|---|
| Control | 180 | 22.79 | 1.907 | 3.126 | 0.6455 | 177 | 0.9029 | 0.001039 |
| Gentamicin | 180 | 26.68 | 1.901 | 3.284 | 0.6423 | 177 | 0.9122 | 0.000760 |
| A1T | 180 | 28/68 | 2.222 | 3.356 | 0.7986 | 177 | 0.9152 | 0.000534 |
| A5T | 180 | 32.07 | 2.329 | 3.468 | 0.8454 | 177 | 0.9240 | 0.000430 |
| A20T | 180 | 38.86 | 1.834 | 3.660 | 0.6064 | 177 | 0.8131 | 0.000930 |
| A40T | 180 | 73.04 | 1.835 | 4.291 | 0.6072 | 177 | 0.8822 | 0.000311 |

* Each frame of the MD simulation was considered as a point.
